# Phenotypic plasticity as a function of genetic polymorphism: thermal dominance reversal in *Drosophila* species with contrasting melanism

**DOI:** 10.1101/2025.05.30.657080

**Authors:** Jean R. David, Béatrice Denis, Pascaline Francelle, Alicia Lemaire, Aparup Das, Sujata Mohanty, Patricia Gibert, Amir Yassin

## Abstract

Phenotypic plasticity is often seen as an alternative adaptive strategy to genetic polymorphism, especially in response to rapid environmental changes. Indeed, a link between plasticity and heterozygosity, *i.e.* the measure of polymorphism, has previously been dismissed. Here, we compare the thermal plasticity of abdominal pigmentation in eight *Drosophila* species, four belonging to the *melanogaster* species group and four to the *montium* group. Despite a conserved developmental pathway for melanin synthesis, the genetic architecture of its variation has significantly evolved, being polygenic in most species (like *D. melanogaster*) and Mendelian or invariable in others. By investigating the thermal plasticity of this trait in species with distinct architectures, we show the degree of plasticity to strongly associate with heterozygosity. Plasticity was resurrected in hybrids between species with no plastic responses but with contrasting melanism, and was higher in heterozygotes in species with simple Mendelian polymorphism. In plastic cases, pigmentation dominance is reversed depending on the developmental temperature. We propose simple genetic models with empirical molecular support to explain this link between phenotypic plasticity and genetic polymorphism. The relationship between these two phenomena, and the impact of each on the evolution of the other, may be more relevant than it is currently appreciated.

## Introduction

Phenotypic plasticity is the ability of a genotype to produce different phenotypes under distinct environmental conditions (Bradshaw 1965; Scheiner 1993; Pigliucci 2001; West-Eberhard 2003; Sultan 2021). When optimal phenotypic values are generated in the different environments, plasticity can facilitate adaptation even in the absence of genetic variation (Ghalambor et al. 2015). This becomes particularly important in cases where the population loses a significant portion of genetic polymorphism, such as after a founder bottleneck during colonization of a new environment, or when the environment change is faster than the rate of accumulation of new adaptive mutations, the so-called ‘buying time’ role of plasticity (Diamond and Martin 2021). Phenotypic plasticity and genetic polymorphism might be considered as distinct alternative adaptive strategies (e.g., Wennersten and Forsman 2012).

Despite the definition of plasticity as a property of a genotype, it is usually studied for complex phenotypic traits whose underlying genotypes are not fully known (e.g., flowering time in *Arabidopsis*, growth in yeasts, etc.) (Goldstein and Ehrenreich 2021; Kovuri et al. 2023). Genotypes do not only differ in the degree of their phenotypic plasticity (e.g., the slope of the reaction norm) but also in their degree of genetic variability, with heterozygous genotypes in non-haploid organisms having more alleles than hemi- or homozygotes. Among the early genetic models of phenotypic plasticity, the Overdominance model explicitly related plasticity to genotypic diversity (Scheiner 1993). The model postulated that plasticity is a function of homozygosity, with the heterozygous genotype considered to be the most advantageous because it is the most robust, *i.e.* the least plastic. This assumption reflected the view of plasticity at the mid-20^th^ century as a developmental nuisance (Lerner 1954; Bradshaw 2006). Indeed, some of the earliest empirical examples of the overdominance model came from studies of wing morphology in *Drosophila melanogaster*, where two mutants showed the highest temperature-sensitivity when found in the homozygous state, but their heterozygote recapitulated the wildtype thermal response, *i.e.* it was less plastic than either mutant (Harnly and Harnly 1936; Schmalhausen 1949). Gillespie and Turelli (1989) extended this view to a multi-locus quantitative genetics model and concluded that stabilizing selection on more environmentally robust heterozygotes will maintain genetic variation in populations. Pigliucci (1992) developed a one-locus, two-allele classical population genetics model for testing the evolution of plasticity under overdominance and concurred with Gillespie and Turelli’s (1989) multilocus result. He concluded that plasticity becomes “increasingly disadvantageous when the system approaches a perfect equilibrium between the probabilities of occurrence of the two environments.”

A direct link between heterozygosity and robustness was not always evident from the empirical literature, and overdominance was disfavored relative to other genetic models such as pleiotropy, *i.e.* plasticity is a function of the expression of a gene, and epistasis, *i.e.* plasticity is a function of interactions between loci. Scheiner (1993) concluded that “plasticity is not a function of heterozygosity”, and Pigliucci (2005), a decade later, stated that “heterozygosity ha[d] little, if anything, to do with the genetic basis of plasticity.” A search using “heterozygo*” in a recent 400-page long book on *Phenotypic Plasticity and Evolution* (Pfennig 2021) yielded no result. However, almost none of the studies cited by Scheiner (1993) and Pigliucci (2005) to reach their conclusions has addressed the link between heterozygosity and plasticity in the most optimal setting similar to Pigliucci’s (1992) theoretical model, *i.e.* comparing the reaction norms of a phenotype with a simple, well-characterized genotypic basis. Conspicuous color polymorphisms are ideally suited for these experiments, because they often involve a simple genetic architecture and color synthesis in many species results from a well conserved gene network (Orteu and Jiggins 2020). In several lineages of moths, simple Mendelian polymorphisms have evolved in response to the industrial revolution, with the melanic wing allele being dominant in some species and recessive in others (Ford 1945). The genetic basis of this industrial melanism has recently been identified to be a conserved micro-RNA close to the locus of the transcription factor *Cortex* (Tian et al. 2024). Remarkably, several butterfly species also show a seasonal color polyphenism, with cold season morphs being darker. van der Burg et al. (2020) showed that mutations at the same locus containing *Cortex*, and consequently the linked micro-RNA, underlie this polyphenism in *Junonia caenia*. Plasticity in simple color polymorphism systems was also observed in *Harmonia* ladybugs morphs (Michie et al. 2010).

Much of our knowledge about the genetic and developmental basis of melanin synthesis comes from studies in *Drosophila*. Pigmentation differences between body segments, sexes, geographical populations, seasons, and species are often due to *cis*- and *trans*-regulatory modifications of melanin synthesis genes, such as *tan (t)*, *yellow (y)* or *ebony (e)*, and their transcription factors, such as *bric-à-brac (bab)*, *POU-domain-motif-3 (pdm3)* or *Abdominal-B (Abd-B)* (Massey and Wittkopp 2016; Hughes et al. 2023). Wild-caught flies from the same locality usually differ in their degree of pigmentation (Figure 1 A; Bastide et al. 2013, 2016; Dembeck et al. 2015; Endler et al. 2016; Castro et al. 2018), but flies collected from cold high altitudes, latitudes, and seasons tend to be dark, even when their progeny is maintained under standard thermal conditions (Telonis-Scott et al. 2011; Bastide et al. 2014, 2016; Takahashi et al. 2014; Negoua et al. 2019; Berardi et al. 2024). Similarly, when full-sib flies issued from the same isofemale line are grown under different temperatures, a negative reaction norm is detected, with flies grown under low temperatures being darker (Figure 1; (Gibert et al. 1996, 1998, 2000; Chakir et al. 2002; Pétavy et al. 2002; Gibert et al. 2004b, 2009; Pétavy et al. 2018; Lafuente et al. 2021, 2024). Thermal plasticity in *D. melanogaster* is due to the higher expression of the *t* and *y* genes in colder temperatures (Gibert et al. 2016, 2017b). A natural variation in the transcription factor *bab* underlying color differences also showed distinct responses to developmental temperature through repression of *t* and *y* in hotter environments (Castro et al. 2018). This parallelism between genetic polymorphism in the wild and phenotypic plasticity in the lab is strongly suggestive for an adaptive role for darker pigmentation in thermoregulation. Indeed, drosophilids with darker cuticle have higher body temperature (Freoa et al. 2023).

**Figure 1.**
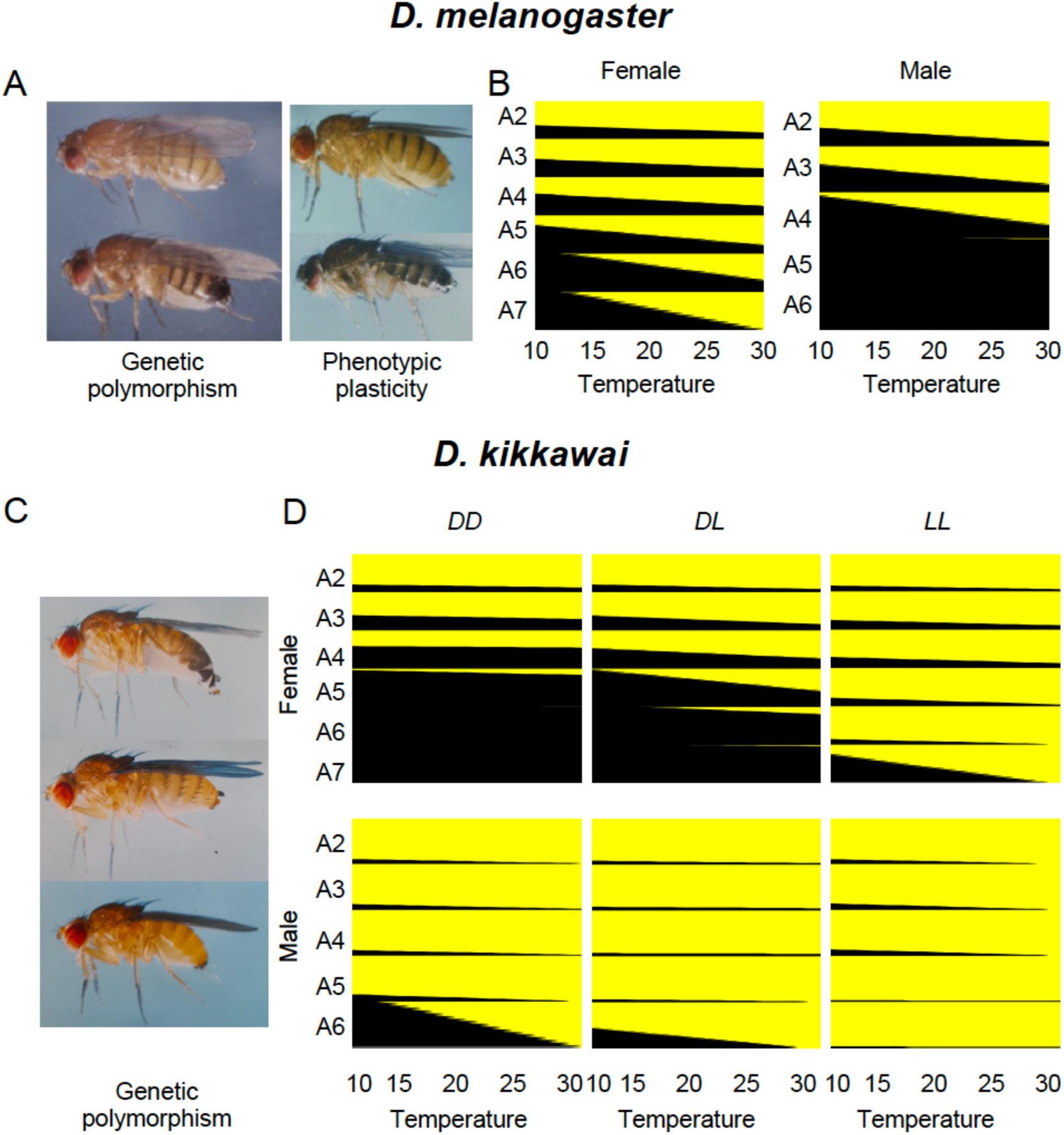
Pigmentation variation in *Drosophila melanogaster* and *D. kikkawai*. A) Variation in abdominal pigmentation in *D. melanogaster* can be due to genetic polymorphism (left) or thermal phenotypic plasticity (right). B) Contour plots showing pigmentation thermal plasticity in *D. melanogaster* females (left) and males (right) at each abdominal segment (A2-A7). C) Variation in abdominal pigmentation in *D. kikkawai* is caused by a single locus with two alleles dark (*D*) and light (*L*) whose effect is more pronounced on females (upper *DD* and middle *LL* photos) than males (lower photo). D) Contour plots showing pigmentation thermal plasticity in *D. kikkawai* females (above) and males (below) for the three genotypes *DD*, *DL*, and *LL*. For each abdominal segment, a reaction norm was inferred as a linear regression slope using mean values available published in Gibert et al. (1999, 2000). Photos are from Jean R. David’s archive.

The genetic architecture of pigmentation variation has dramatically evolved between drosophilid species. Whereas female abdominal pigmentation variation is continuous and polygenic in *D. melanogaster* as well as most species, it is discrete and monogenic in multiple species (Payant 1986; Yassin et al. 2016a,b). Female-limited Mendelian melanism has evolved in *D. erecta* of the *melanogaster* group and in 22 species of the *montium* group belonging to five distinct subgroups (Yassin et al. 2016b; Yassin 2018). Of the 16 *montium* species where the genetics was studied, the dark allele was dominant in 13 species (Yassin et al. 2016b; Prigent et al. 2020). The phenotypic plasticity of abdominal pigmentation was investigated by Gibert et al. (1999) in one species, namely *D. kikkawai*, where females have two morphs and males have a light abdomen. The authors found major effect of genotype, segment, sex, and temperature (Figure 1B). In some other species, pigmentation is nearly fixed with very slight variability even between the sexes. This is the case of the two sister species, *D. santomea* and *D. yakuba*, where both males and females’ posterior abdomens are invariably light or dark, respectively (Lachaise et al. 2000; Jeong et al. 2008; David et al. 2022).

In this paper, we compared the level of pigmentation phenotypic plasticity in eight species with distinct genetic architectures grown under five thermal regimes. To this end, we focused on the effect of heterozygosity, as it can be seen in heterozygous genotypes of five species with monogenic architectures as well as in species hybrids (*i.e.* the ultimate viable heterozygotes). Our results support a positive link between plasticity and the degree of genetic polymorphism that was not previously assumed.

## Materials and Methods

### Strains used and fly husbandry

We studied five cases where for each the thermal phenotypic plasticity can be compared between homozygous dark and light lines and their reciprocal hybrids. Four cases considered within-species monogenic Mendelian color polymorphism with one (*D. erecta*) from the *melanogaster* species group and three cases (*D. jambulina*, *D. burlai*, and *D.* cf. *bocqueti*) from its sister clade, the *montium* species group. For each of these species homozygous dark and light lines for the color causing allele were established through selection from a polymorphic mass culture (Table 1). The monogenic basis of female pigmentation in these species are known (Yassin et al. 2016a,b; Prigent et al. 2020). In two species (*D. erecta* and *D. burlai*), the dark allele is dominant, although the causative locus is different being the X-linked *tan* in *D. erecta* and the autosomal *pdm3* in *D. burlai* (Yassin et al. 2016a,b). In the two other species (*D. jambulina* and *D.* cf. *bocqueti*), the light allele is dominant (Watanabe et al. 1982; Prigent et al. 2020), but the causative locus is still not mapped although color variation associates with polymorphism in *pdm3* in the two species (Fukutomi et al. 2025). The fifth case involved hybridization between the two sister species *D. santomea* and *D. yakuba* of the *melanogaster* group, whose posterior segments are either monomorphically light or dark, respectively (Table 1). The pigmentation difference between these two species is controlled by mutations in at least six loci (Liu et al. 2019; David et al. 2022). All strains were kept in a rearing room at 21°C with 12:12 dark:light cycles on a food medium composed of 10 g agar, 101 g sugar, 25 g yeast, 100 g corn flour, 6.9 g nipagin, and 50 mL ethanol per 1 L. We also obtained published data of the thermal plasticity response of *D. melanogaster* (Gibert et al. 2000), the nomitypical species of the *melanogaster* group with polygenic basis of pigmentation variation, and the monogenic *D. kikkawai* (Gibert et al. 1999) for comparison purposes (Table 1).

**Table 1.**
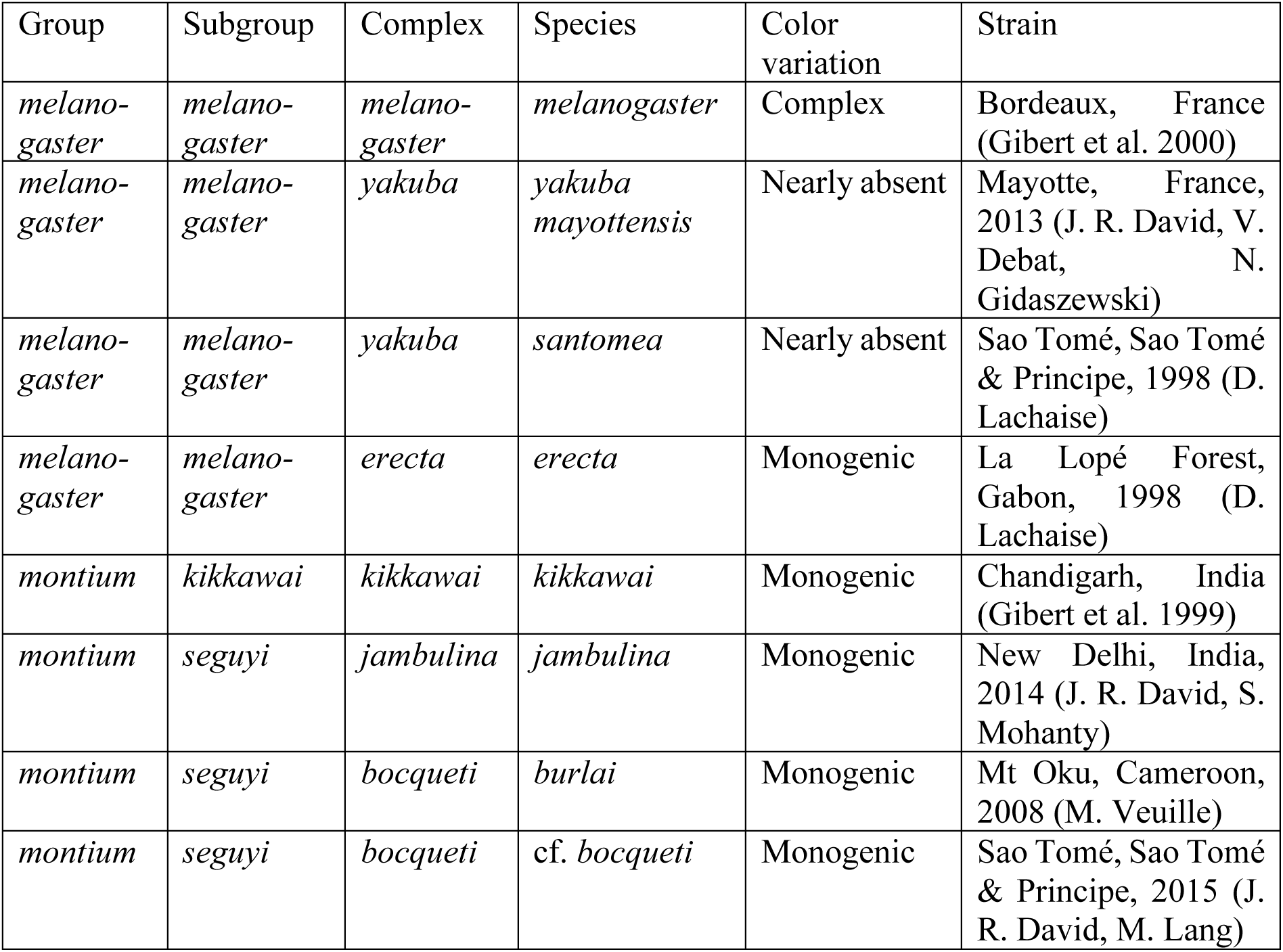
*Drosophila* strains analyzed in this study.

| Group | Subgroup | Complex | Species | Color variation | Strain |
| --- | --- | --- | --- | --- | --- |
| *melano-gaster* | *melano-gaster* | *melano-gaster* | *melanogaster* | Complex | Bordeaux, France (Gibert et al. 2000) |
| *melano-gaster* | *melano-gaster* | *yakuba* | *yakuba mayottensis* | Nearly absent | Mayotte, France, 2013 (J. R. David, V. Debat, N. Gidaszewski) |
| *melano-gaster* | *melano-gaster* | *yakuba* | *santomea* | Nearly absent | Sao Tomé, Sao Tomé & Príncipe, 1998 (D. Lachaise) |
| *melano-gaster* | *melano-gaster* | *erecta* | *erecta* | Monogenic | La Lopé Forest, Gabon, 1998 (D. Lachaise) |
| *montium* | *kikkawai* | *kikkawai* | *kikkawai* | Monogenic | Chandigarh, India (Gibert et al. 1999) |
| *montium* | *seguyi* | *jambulina* | *jambulina* | Monogenic | New Delhi, India, 2014 (J. R. David, S. Mohanty) |
| *montium* | *seguyi* | *bocqueti* | *burlai* | Monogenic | Mt Oku, Cameroon, 2008 (M. Veuille) |
| *montium* | *seguyi* | *bocqueti* | *cf. bocqueti* | Monogenic | Sao Tomé, Sao Tomé & Príncipe, 2015 (J. R. David, M. Lang) |

### Pigmentation scoring

For monogenic species, we followed the nomenclature given in Gibert et al.’s (1999) paper, referring to the light and dark alleles as *L* and *D*, respectively, and to the three genotypes as *LL*, *LD* or *DL*, and *DD*. For heterozygotes, the allele from the mother is given first depending on the cross. For each genotype, pigmentation was scored on >4 days old 10 males and 10 females as in Yassin et al. (2022) following the pigmentation score scale proposed by David et al. (1990). This pigmentation score visually quantifies the proportion of the pigmented part of an abdominal segment when viewed on the lateral side of a fly under a stereoscope and it goes from 0 (no pigmentation) to 10 (fully pigmented). It is therefore different from other pigmentation scores in *Drosophila* that depend on the intensity of melanism or the length of the pigmented stripe quantified after numerical photographing (e.g., Bastide et al. 2014; Lafuente et al. 2021). Scorings were made by the same observer (the first author) between 2013 and 2019 (but mostly between March 2016 and April 2017) to limit observation bias. Data were manually entered on a printed sheet with a template table for morphometric data and then the means for each segment were calculated in a separate sheet. All sheets are archived. Scorings were made by the same observer to limit observation bias between 2013 and 2019 (but mostly between March 2016 and April 2017). Abdomens of some flies at extreme temperatures (see below) were dissected and mounted on microscopic slides and photomicrographed (cf. data associated with this study). In each case, pigmentation was scored on six and five abdominal segments in females and males respectively (denoted A2-7 with F and M for females and males, respectively).

Flies were grown at five temperatures going from 12°C to 31°C with 3-4° increments. For most cases, the five categories were 14, 17, 21, 25, and 28°C, but a few cases were scored at 19 (*D. jambulina DD*), 29 (*D.* cf. *bocqueti DL*), and 30°C (*D. erecta DD*, *DL*, *LD*, and *LL*). Mortality during development was not assessed. For *D. melanogaster* and *D. kikkawai*, only mean values per genotype were retrieved from the literature for seven (12, 14, 17, 21, 25, 28, and 31°C) and four (12, 17, 25, and 30°C) temperature degrees, respectively (Gibert et al. 1999, 2000). For three genotypes < 10 flies per sex were available at the time of observation (namely, *D.* cf. *bocqueti LL* males at 21°C (*N* = 2), *D. jambulina DD* females and males at 25°C (*N* = 5). Consequently, temperature was coded either as a degree or as a class depending on the statistical test used (see below), with the five classes being: cold (12-14°C), mildly cold (17-19°C), mild (21°C), mildly hot (25°C), and hot (28-31°C). In total, pigmentation was scored for nearly 8,703 abdominal segments over 1,582 flies for the four species with monogenic melanism. For the hybridization experiment between *D. santomea* and *D. yakuba*, only the sheet recording the means for each genotype per sex per temperature was retrieved for the five temperatures. However, photomicrographs of abdominal segments for the four genotypes (parental strains and reciprocal crosses) at three temperatures (14, 21, and 28°C) are provided in the data associated with this study. Analyses on the whole set of temperatures for this experiment were therefore limited to the mean values.

### Statistical analyses

All statistical analyses were conducted using R (R Core Team 2023). As a serially homologous trait, abdominal pigmentation can be tested either as in an integrative (*i.e.* the overall pigment production) or a modular way (*i.e.* pigmentation patterns across segments). Indeed, pigmentation in all segments is produced by the same melanin synthesis genes and their variants that each individual carries. Previous works in *D. melanogaster* and *D. kikkawai* have shown a substantial degree of correlation especially among adjacent segments (David et al. 1990; Gibert et al. 1999, 2000). However, pigmentation variation associates with different genetic variants in different abdominal segment in *D. melanogaster* measured on the same isogenic lines (Dembeck et al. 2015), and the reaction norms of the thermal plasticity are different even among adjacent abdominal segments in *D. simulans* (Gibert et al. 1998). To test for the independence between the abdominal segments we conducted correlation analyses on the whole individual-based dataset using the Corrplot package in R (Wei and Simko 2024), and found indeed substantial degrees of correlations which were stronger among adjacent segments (see Results). Consequently, we conducted our further analyses either on each segment separately or using the sum of all segments per an individual depending on the test.

We first conducted a multi-factor Analysis of Variance (ANOVA) on the individual-based dataset for the four species with monogenic melanism to test of the effect of each factor (species, genotype, sex, segment, and temperature) as well as of their interactions on pigmentation variation. We estimated the effect of these factors as the proportion of variance explained (*η*^2^) using the effectsize package (Ben-Shachar et al. 2020) in R. To test for the effect of reciprocal crosses, we conducted the same analysis only on the *DL* and *LD* genotypes. Because no significant difference was found between these genotypes, they were pooled into a single F_1_ heterozygous category for subsequent analyses. For a balanced ANOVA, equal number of segments were used for each sex. To do so in *D. kikkawai*, Gibert et al. (1999) excluded the most posterior female abdominal segment, *i.e.* FA7. However, subsequent analyses showed that this segment is among the most variable within and between Drosophila species (Bastide et al. 2013; Yassin et al. 2016a,b). Consequently, we excluded here the most anterior female segment, *i.e.* A2, which usually shows less variation and strong correlation with other anterior segments (see Results). Similarly, temperature classes rather than temperature degrees were used to test the effect of temperature in a balanced way across species and genotypes.

Previous analyses have shown the shape of the thermal reaction norms to differ between segments of the same isofemale lines. For example, segments 5, 6, and 7 in *D. simulans* females have quadratic, linear, and sigmoid (logistic) norms, respectively, although all show the same response of higher melanism in colder environments (Gibert et al. 1998). Because our main focus in this paper is to compare the level of plasticity between heterozygotes and homozygotes rather than the diversity of the shape of the reaction norms, linear regressions of pigmentation scores over temperature degrees were conducted using the lm function in R for every segment as well as for the sum of all segments, and the slope of the regression line (*b*) was considered as a proxy for the degree of plasticity. Regression analyses were conducted on the whole individual-based dataset rather than on averages for the measurements generated by this study, although for the data retrieved from the literature for *D. melanogaster* and *D. kikkawai*, and for the hybridization experiment, only averages were available.

To visualize the patterns of segment-specific diversity for each genotype per sex per species we used a customized perl script to translate the linear regression equations generated above into a pigmentation pattern. Each segment was divided into a grid of 100 centiles so that a pigmentation score of 2.2 for a segment means that the 22 most posterior centiles of that segment have a pigmentation score of 10, whereas the 78 most anterior centiles have a score of 0. A 2D contour map was then drawn using the Plotly package in R (Sievert 2020). For the overall pigmentation score (*i.e.* sum of all segments), 2D plots were drawn for each sex per species using the plot function in R. Genetic dominance was inferred as the difference between the slope of pooled heterozygotes (*DL* and *LD*) and the slope of the pooled homozygotes (*DD* and *LL*), with dominance reversal being witnessed at the intersection between the two slopes within the range of vital temperatures between 10 and 30°C. To test if heterozygotes had different plasticity than homozygotes, we ranked the slope of each of the three genotypes from the highest to the lowest as 3, 2, and 1 for each species and conducted a non-parametric Wilcoxon test on the relation between rank and genotype.

For the hybridization experiment between *D. santomea* and *D. yakuba*, only mean scores for each genotype per sex per temperature were available (see above). We first conducted a multi-factor ANOVA on these values, after excluding female A2 and considering the five temperature classes as for the species with the monogenic melanism. We also conducted ANOVA on only the hybrid genotypes to test for the effect of reciprocal crosses on the whole dataset. To quantify and visualize plasticity at each segment as well as on the total pigmentation scores we conducted linear regression analyses on the mean values as above. We note that for species with monogenic melanism, regression slopes strongly correlated between the whole datasets and the means (*r* = 0.997, *P* < 2.2 x 10^-16^; Student’s t *P* = 0.871), although the standard error of the slope value was significantly reduced in individual-based analyses (*r* = 0.849, *P* < 2.2 x 10^-16^; Student’s t *P* = 1.671 x 10^21^). Consequently, the slope reported for the mean values of this experiment correctly reflects the plasticity response.

## Results

### Pigmentation thermal plasticity varies with species, sex, segment, and genotype in species with monogenic melanism

ANOVA on the whole dataset of the four species with monogenic melanism indicated a strong effect of all factors and their interactions (Table 2). However, the most significant effect was due to segment position (explaining 30% of the variance), followed by temperature and genotype (5%), then species (2%) and sex (1%). To test for the effect of reciprocal crosses, we conducted the same ANOVA on the data set of only heterozygous genotypes (*LD* and *DL*). We found that the genotype had no effect at all, whether individually or in interaction with other factors. For simplicity, we pooled data for heterozygous individuals issued from reciprocal crosses into a single genotypic group (F_1_). When we conduct the analysis using the complete genotypic dataset (*i.e.* with pooled *LD* and *DL*), the ANOVA results showed the significant effect of all factors and their interactions.

**Table 2.** Analysis of variance (ANOVA) for pigmentation variation in four species with monogenic melanism. *D* and *L* refers to dark and light alleles, respectively. Df = degree of freedom, GEN = genotype, SEG = segment position, SPE = species, and TCL = temperature class. *** *P* < 0.001.

| Genotypes | *DD, DL, LD, LL* |  |  | *DL, LD* |  |  | *DD, F<sub>1</sub>, LL* |  |  |
| --- | --- | --- | --- | --- | --- | --- | --- | --- | --- |
| Factor | Df | F | $\eta^2$ | Df | F | Eta2 | Df | F | $\eta^2$ |
| SPE | 3 | 222.438*** | 0.02 | 3 | 104.487*** | 0.02 | 3 | 222.897*** | 0.02 |
| GEN | 3 | 546.944*** | 0.05 | 1 | 0.433 | <0.01 | 2 | 821.871*** | 0.05 |
| TCL | 1 | 1459.131**<br>* | 0.05 | 1 | 1183.554**<br>* | 0.08 | 1 | 1462.142**<br>* | 0.05 |
| SEX | 1 | 337.125*** | 0.01 | 1 | 87.979*** | <0.01 | 1 | 337.820*** | 0.01 |
| SEG | 1 | 9325.489**<br>* | 0.30 | 1 | 5121.901**<br>* | 0.36 | 1 | 9344.735**<br>* | 0.30 |
| SPE:GEN | 9 | 21.791*** | <0.01 | 3 | 0.328 | <0.01 | 6 | 32.573*** | <0.01 |
| SPE:TCL | 3 | 70.243*** | <0.01 | 3 | 79.879*** | 0.02 | 3 | 70.388*** | <0.01 |
| GEN:TCL | 3 | 55.130*** | <0.01 | 1 | 3.112 | <0.01 | 2 | 81.154*** | <0.01 |
| SPE:SEX | 3 | 156.685*** | 0.02 | 3 | 115.028*** | 0.02 | 3 | 157.008*** | 0.02 |
| GEN:SEX | 3 | 306.050*** | 0.03 | 1 | 0.107 | <0.0<br>1 | 2 | 459.963*** | 0.03 |
| TCL:SEX | 1 | 96.736*** | <0.0<br>1 | 1 | 178.999*** | 0.01 | 1 | 96.936*** | <0.0<br>1 |
| SPE:SEG | 3 | 736.924*** | 0.07 | 3 | 503.505*** | 0.11 | 3 | 738.445*** | 0.07 |
| GEN:SEG | 3 | 518.281*** | 0.05 | 1 | 0.781 | <0.0<br>1 | 2 | 778.596*** | 0.05 |
| TCL:SEG | 1 | 45.414*** | <0.0<br>1 | 1 | 119.222*** | <0.0<br>1 | 1 | 45.508*** | <0.0<br>1 |
| SEX:SEG | 1 | 1118.545**<br>* | 0.04 | 1 | 308.347*** | 0.02 | 1 | 1120.853**<br>* | 0.04 |
| SPE:GEN:TCL | 9 | 11.695*** | <0.0<br>1 | 3 | 1.447 | <0.0<br>1 | 6 | 16.783*** | <0.0<br>1 |
| SPE:GEN:SEX | 9 | 13.581*** | <0.0<br>1 | 3 | 0.138 | <0.0<br>1 | 6 | 20.337*** | <0.0<br>1 |
| SPE:TCL:SEX | 3 | 20.594*** | <0.0<br>1 | 3 | 32.869*** | <0.0<br>1 | 3 | 20.636*** | <0.0<br>1 |
| GEN:TCL:SEX | 3 | 34.231*** | <0.0<br>1 | 1 | 0.056 | <0.0<br>1 | 2 | 51.421*** | <0.0<br>1 |
| SPE:GEN:SEG | 9 | 22.254*** | <0.0<br>1 | 3 | 0.051 | <0.0<br>1 | 6 | 33.421*** | <0.0<br>1 |
| SPE:TCL:SEG | 3 | 10.443*** | <0.0<br>1 | 3 | 26.690*** | <0.0<br>1 | 3 | 10.464*** | <0.0<br>1 |
| GEN:TCL:SEG | 3 | 30.370*** | <0.0<br>1 | 1 | 0.079 | <0.0<br>1 | 2 | 45.606*** | <0.0<br>1 |
| SPE:SEX:SEG | 3 | 55.643*** | <0.0<br>1 | 3 | 96.490*** | 0.02 | 3 | 55.758*** | <0.0<br>1 |
| GEN:SEX:SEG | 3 | 347.385*** | 0.03 | 1 | 1.411 | <0.0<br>1 | 2 | 521.377*** | 0.03 |
| TCL:SEX:SEG | 1 | 115.968*** | <0.0<br>1 | 1 | 227.193*** | 0.02 | 1 | 116.208*** | <0.0<br>1 |
| SPE:GEN:TCL:SEX | 9 | 6.391*** | <0.0<br>1 | 3 | 0.085 | <0.0<br>1 | 6 | 9.559*** | <0.0<br>1 |
| SPE:GEN:TCL:SEG | 9 | 9.518*** | <0.0<br>1 | 3 | 0.412 | <0.0<br>1 | 6 | 14.079*** | <0.0<br>1 |
| SPE:GEN:SEX:SEG | 9 | 20.399*** | <0.0<br>1 | 3 | 0.212 | <0.0<br>1 | 6 | 30.545*** | <0.0<br>1 |
| SPE:TCL:SEX:SEG | 3 | 25.856*** | <0.0<br>1 | 3 | 49.527*** | 0.01 | 3 | 25.910*** | <0.0<br>1 |
| GEN:TCL:SEX:SEG | 3 | 45.620*** | <0.0<br>1 | 1 | 0.207 | <0.0<br>1 | 2 | 68.457*** | <0.0<br>1 |
| SPE:GEN:TCL:SEX:SEG | 9 | 11.697*** | <0.0<br>1 | 3 | 0.097 | <0.0<br>1 | 6 | 17.529*** | <0.0<br>1 |
| Residuals | 778<br>0 |  |  | 393<br>6 |  |  | 781<br>2 |  |  |

### Higher plasticity of overall pigmentation in heterozygous genotypes in species with monogenic melanism

Despite the strong effect of body segmentation on pigmentation diversity between species, sexes, and genotypes, previous studies have reported strong correlation in pigmentation between adjacent segments (David et al. 1990; Gibert et al. 1999, 2000). We tested this hypothesis using individual data for the four species and indeed found substantial degrees of correlations, especially among adjacent segments, indicating the non-independence of segment-based effect (Figure 2A). Consequently, to exclude such non-independence, we summarized pigmentation variation as sum scores across segments per individual. A multi-factor ANOVA on the sum pigmentation scores showed a significant effect of all factors and their interactions, with the strongest effect being of temperature (explaining 21% of the variance), followed by genotype (18%) and species (11%). To visualize the effect of these three factors, we plotted sum pigmentation scores of each genotype per species over temperature. In all cases, a significant negative linear relation was found, with pigmentation increases in colder environments (Figure 2B). Adjusted R^2^ ranged from 0.339 to 0.899, indicating that some relationships may better be explained by more complex non-linear regressions. However, for the sake of comparisons across species and genotypes, we used the slope of the linear regression of each genotype as a proxy for the degree of phenotypic plasticity. Species greatly differed in these slopes (averaged over three genotypes and two sexes), being highest in *D. jambulina* (mean *b*=-1.189±0.617), followed by *D.* cf. *bocqueti* (*b*=-0.781±0.329), *D. burlai* (*b*=-0.470±0.010), and *D. erecta* (*b*=-0.441±0.080). For each species, we ranked the three genotypes from the most plastic (*i.e.* strongest slope) to the least plastic, and found that heterozygosity strongly correlated with higher plasticity (Wilcoxon’s W = 16, *P* = 0.002).

**Figure 2.**
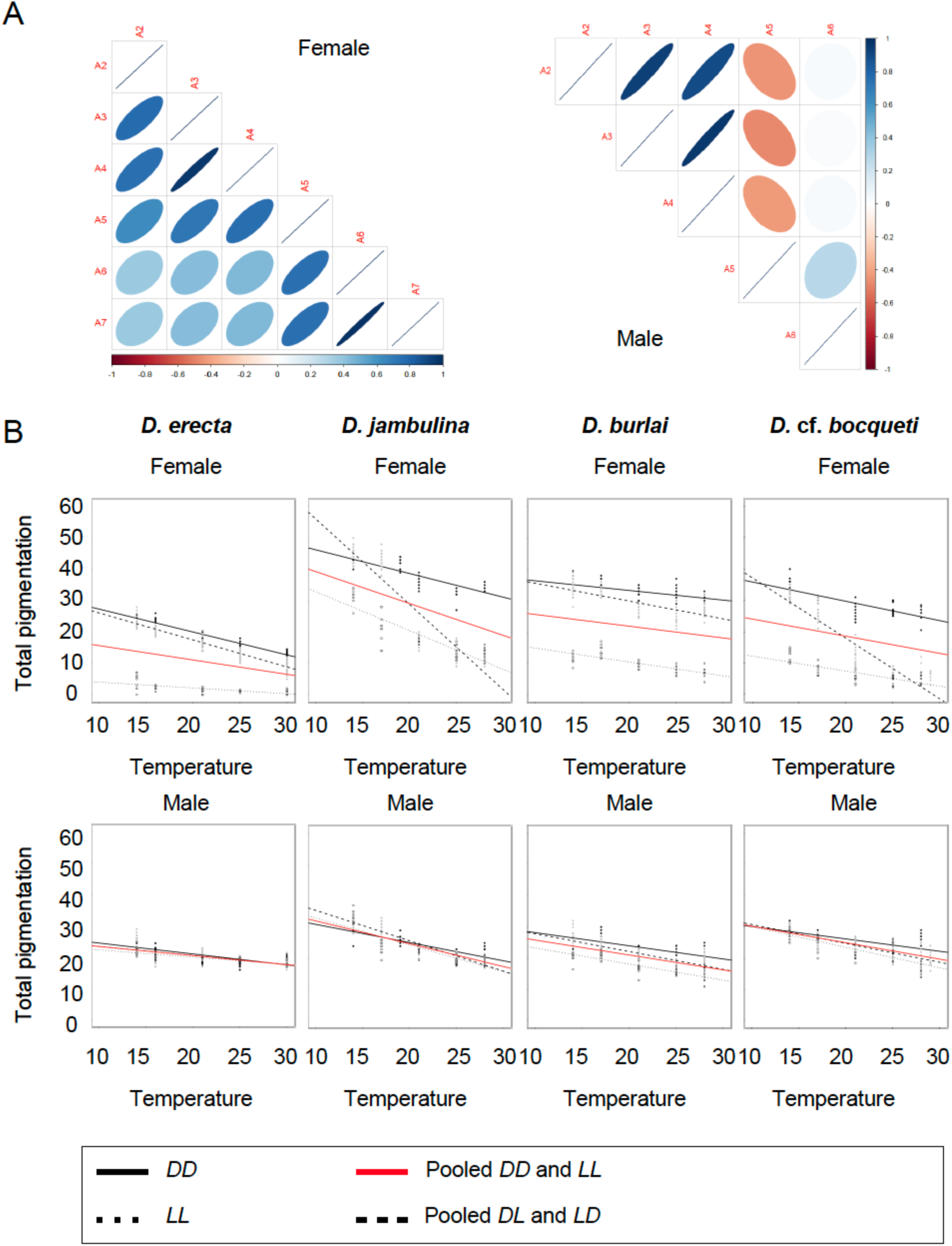
Correlation between abdominal segments and plasticity of total pigmentation in species with monogenic melanism. A) Correlation between pigmentation scores of abdominal segment in females (left) and males (right) showing stronger correlation between adjacent segments. B) Linear regression of total pigmentation (sum of segment scores) over developmental temperature in homozygous dark (*DD*) and light (*LL*) genotypes and pooled heterozygotes (*DD* and *LL*). Dominance can be inferred as the difference between the slope of the pooled heterozygotes (black dashed) and that of the pooled homozygotes (red solid).

### Higher plasticity of heterozygotes associates with dominance reversal

The stronger slope of the heterozygote relative to the homozygotes indicate that dominance between the dark *D* and light *L* alleles likely changes with developmental temperature. To evaluate this effect, we inferred the slope of pooled homozygous genotypes, which is a proxy of the mid-parent values across temperatures. We found indeed that the slope of the heterozygotes approaches this mid-parent slope in almost all cases at higher temperature, indicating a reduction of the dominance of the *D* allele with increasing temperature. The heterozygotes’ slope intercrossed that of the mid-parent at 17°C in *D.* cf. *bocqueti* males, 19°C in *D.* cf. *bocqueti* females, 20°C in *D. jambulina* females, 25°C in *D. jambulina* males and 30°C in *D. burlai* males, indicating a reversal of dominance within the vital temperature range between 10 and 30°C. In two cases, *D. erecta* and *D. burlai* females the intercrossing was expected at >30°C, *i.e.* the *D* allele is dominant throughout the vital range although with decreasing dominance. In *D. erecta* males, the slope superimposed that of the mid-parent throughout the temperature range, indicating a non-plastic codominance of the *D* and *L* alleles. Visual depiction of the regression slope at each segment showed differences between the segments especially towards the posterior of the abdomen (Figures 3 and 4). The reversal of dominance in *Drosophila jambulina* is interesting, since the dominance relationship in this species is ambiguous. Parkash and Sharma (1978) found dark dominance for an Indian population, whereas Watanabe et al. (1982) found light dominance also for Indian populations and concluded that dominance in *D. jambulina* was “a great problem existing in a simple genetic phenomenon”. Our results suggest that Parkash and Sharma (1978) and Watanabe et al. (1982) might have conducted their experiments at different temperatures, hence reaching contradictory results. For *D.* cf. *bocqueti*, Prigent et al. (2020) reported that the light allele was dominant in the Sao Tomé strains that we analyze here. Our results indicate that this statement is temperature-dependent. However, because both *D. jambulina* and *D.* cf. *bocqueti*, are tropical, predominantly living in environments with an average temperature ≥21°C, the light allele may be considered the dominant one in nature.

**Figure 3.**
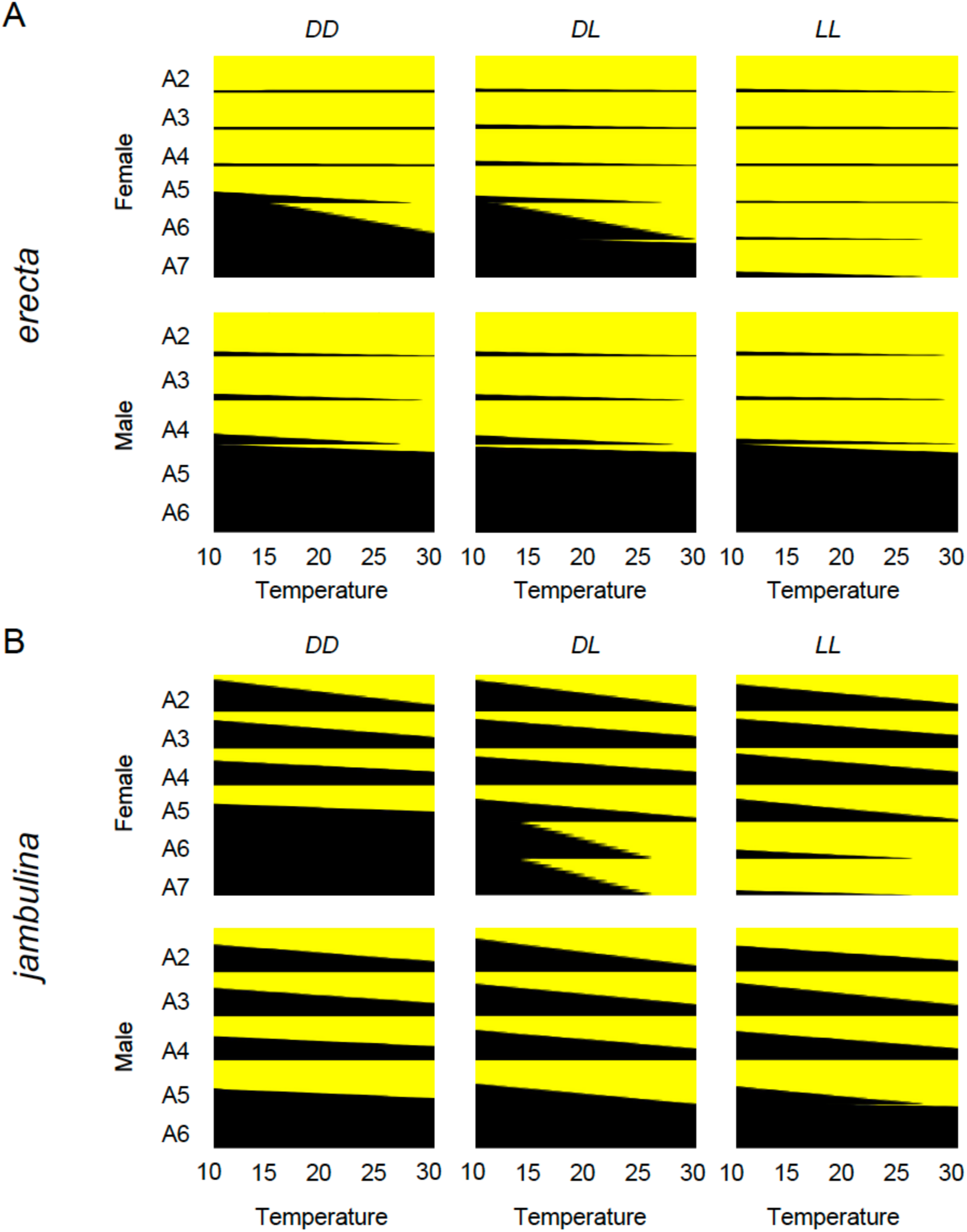
Thermal phenotypic plasticity of pigmentation on successive abdominal segments in *D. erecta* (A) and *D. jambulina* (B). For each panel, contour plots show pigmentation thermal plasticity (proportion of black stripe) in females (above) and males (below) for the three genotypes *DD*, *DL*, and *LL*. For each abdominal segment, a reaction norm was inferred as a linear regression slope using individual values.

**Figure 4.**
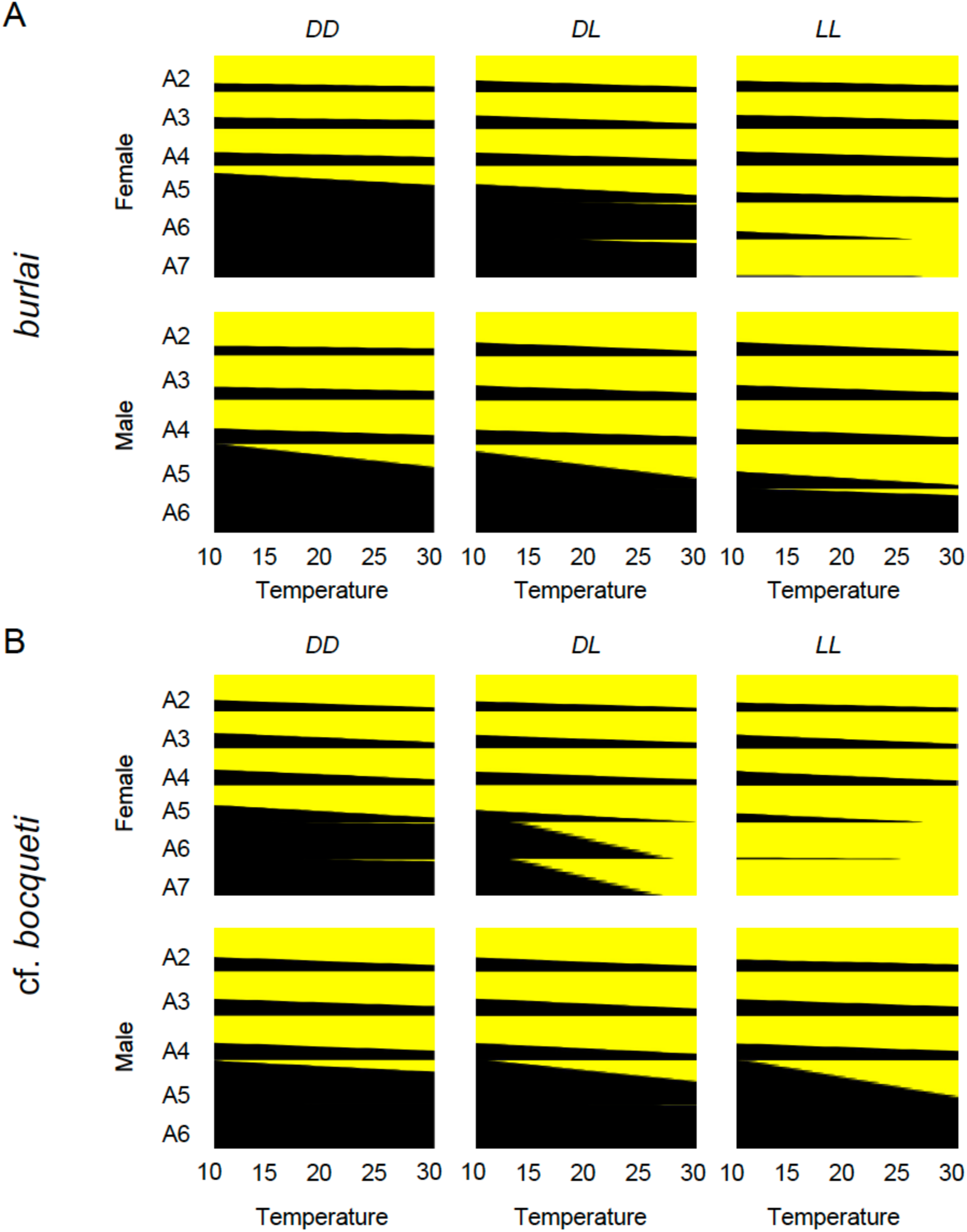
Thermal phenotypic plasticity of pigmentation on successive abdominal segments in *D. burlai* (A) and *D.* cf. *bocqueti* (B). For each panel, contour plots show pigmentation thermal plasticity (proportion of black stripe) in females (above) and males (below) for the three genotypes *DD*, *DL*, and *LL*. For each abdominal segment, a reaction norm was inferred as a linear regression slope using individual values.

### Plasticity increases in hybrids of D. santomea and D. yakuba

Multi-factor ANOVA showed all factors but sex to have a significant effect on abdominal pigmentation in *D. santomea*, *D. yakuba*, and their hybrids, with the genotype being the most important factor (accounting for nearly 39% of the variance), followed by the segment position (25%), and temperature (4%) (Table 3). Indeed, the two parental species are known to show very little sexual dimorphism, but very different pigmentation especially on the posterior segments, with *D. yakuba* and *D. santomea* being nearly completely dark and light, respectively (David et al. 2022; Yassin et al. 2022). Contrary to species with monogenic melanism, multi-factor ANOVA on the heterozygotes from the reciprocal crosses showed a significant effect of the genotype (Table 3).

**Table 3.** Analysis of variance (ANOVA) for pigmentation variation in *D. santomea* (*LL*), *D. yakuba* (*DD*), and their hybrids. *D* and *L* refers to dark (*D. yakuba*) and light (*D. santomea*) alleles, respectively. Df = degree of freedom, GEN = genotype, SEG = segment position, and TCL = temperature class. ** *P* < 0.01; *** *P* < 0.001.

| Experiment | *DD, DL, LD, LL* |  |  | *DL, LD* |  |  |
| --- | --- | --- | --- | --- | --- | --- |
| Factor | Df | F | $\eta^2$ | Df | F | $\eta^2$ |
| GEN | 3 | 230.863*** | 0.39 | 1 | 43.272*** | 0.08 |
| TCL | 1 | 64.244*** | 0.04 | 1 | 65.130*** | 0.12 |
| SEX | 1 | 2.511 | 1.41e-03 | 1 | 3.352. | 5.97e-03 |
| SEG | 1 | 450.899*** | 0.25 | 1 | 238.814*** | 0.43 |
| GEN:TCL | 3 | 7.945*** | 0.01 | 1 | 0.132 | 2.35e-04 |
| GEN:SEX | 3 | 10.601*** | 0.02 | 1 | 23.579*** | 0.04 |
| TCL:SEX | 1 | 0.001 | 3.04e-07 | 1 | 0.321 | 5.71e-04 |
| GEN:SEG | 3 | 73.875*** | 0.12 | 1 | 16.177*** | 0.03 |
| TCL:SEG | 1 | 22.129*** | 0.01 | 1 | 35.368*** | 0.06 |
| SEX:SEG | 1 | 13.212*** | 7.42e-03 | 1 | 14.030*** | 0.02 |
| GEN:TCL:SEX | 3 | 0.683 | 1.15e-03 | 1 | 0.970 | 1.73e-03 |
| GEN:TCL:SEG | 3 | 7.495*** | 0.01 | 1 | 0.835 | 1.49e-03 |
| GEN:SEX:SEG | 3 | 5.626** | 9.48e-03 | 1 | 9.489** | 0.02 |
| TCL:SEX:SEG | 1 | 2.662 | 1.50e-03 | 1 | 3.439. | 6.13e-03 |
| GEN:TCL :SEX:SEG | 3 | 1.686 | 2.84e-03 | 1 | 2.550 | 4.54e-03 |
| Residuals | 208 |  |  | 104 |  |  |

Total pigmentation (summed over all segments) showed significant thermal plasticity in both sexes in *D. yakuba* (females: *b* = -0.365 ± 0.039, *P* < 0.001; males: *b* = -0.513 ± 0.057, *P* < 0.001) but only and to a lesser degree in *D. santomea* females (females: *b* = -0.093 ± 0.020, *P* < 0.01 males: *b* = -0.145 ± 0.057, *P* = 0.065) (Figure 5). The slope of the linear regression was higher in hybrids, approaching -1 in females (*DL*: *b* = -1.277 ± 0.155, *P* < 0.01; *LD*: *b* = -0.941 ± 0.092, *P* < 0.001) and males (*DL*: *b* = -0.866 ± 0.161, *P* < 0.01; *LD*: *b* = -1.020 ± 0.096, *P* < 0.001) (Figure 5). As for species with monogenic melanism, the slope of the heterozygotes approached or crossed that of the mid-parents indicating a reversal of dominance. In females of reciprocal crosses, the intercrossing occurred at intermediated temperatures (17-20°C). In males, on the other hand, the slope of the heterozygotes approached that of the pooled homozygotes but in opposite sides. For the *san x yak (LD)* males, the *santomea* light phenotype was dominant throughout the range, approaching codominance at cold temperatures. For the *yak x san (DL)* males, the opposite was found with the *yakuba* dark phenotype was dominant throughout the range approaching codominance at hot temperatures. This indicates a strong effect of the X chromosome, which carries two melanin-synthesis genes with loss-of-function mutations in *D. santomea* (Liu et al. 2019; David et al. 2022; see Discussion).

**Figure 5.**
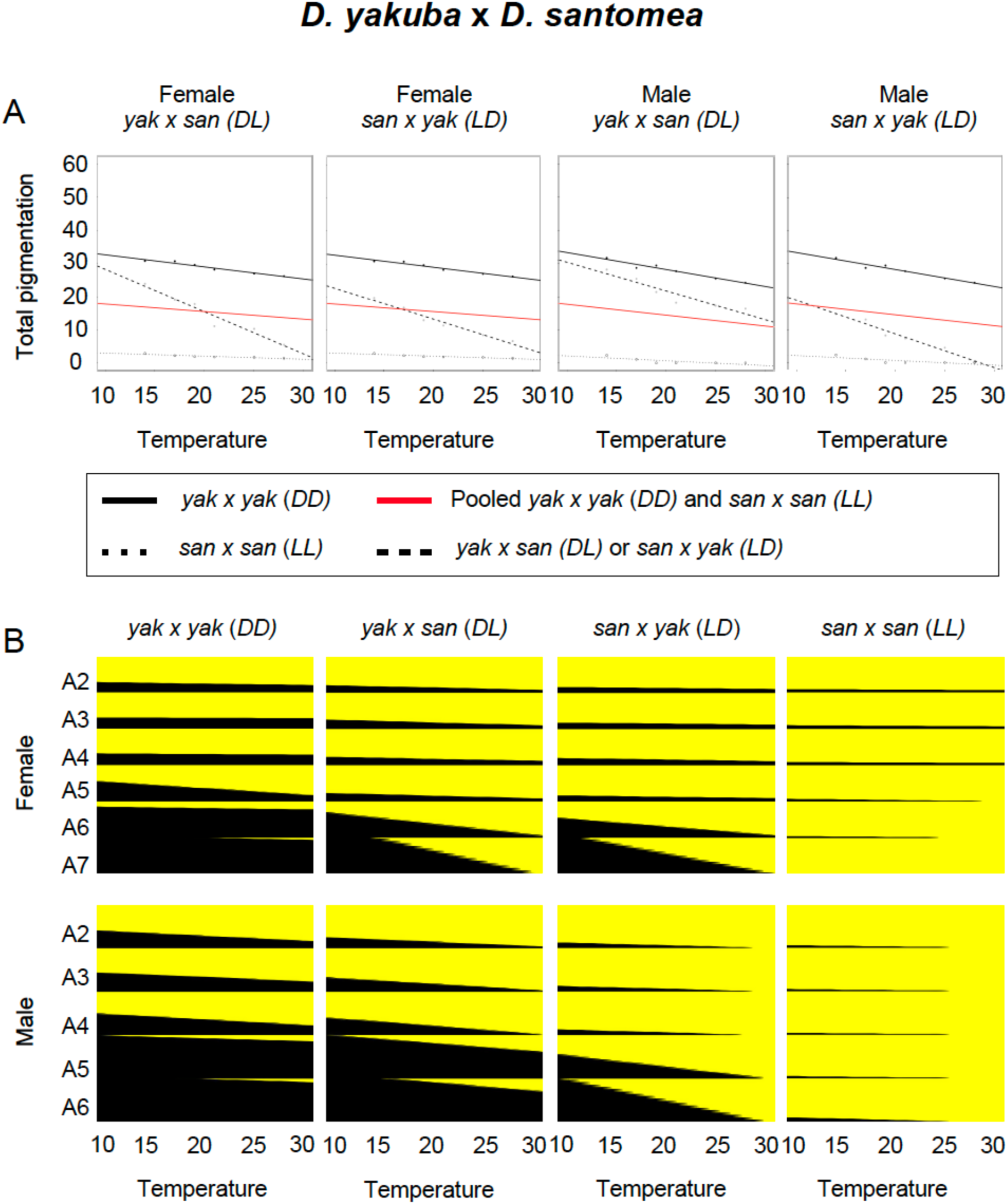
Thermal phenotypic plasticity of abdominal pigmentation in *D. yakuba*, *D. santomea*, and their hybrids. A) Linear regression of total pigmentation (sum of segment scores) over developmental temperature in homozygous dark (*DD*) and light (*LL*) genotypes and pooled heterozygotes (*DD* and *LL*). Dominance can be inferred as the difference between the slope of the pooled heterozygotes (black dashed) and that of the pooled homozygotes (red solid). B) Plasticity patterns at each abdominal segment. For each panel, contour plots show pigmentation thermal plasticity (proportion of black stripe) in females (above) and males (below) for the three genotypes *DD*, *DL*, and *LL*. For each abdominal segment, a reaction norm was inferred as a linear regression slope using mean values (see text).

As indicated from the ANOVA analyses, segment position had very important effect on pigmentation, especially in comparing reciprocal crosses where it explained 43% of the variance (Table 3). This is perfectly seen in the visual depiction for each segment, where differences in pigmentation plasticity were more pronounced in posterior segments (Figure 5).

## Discussion

### The molecular basis of the association between plasticity and heterozygosity in Drosophila

Although it has long been recognized that genetic variation of plasticity exists, the nature of this variation remains elusive. Early works suggested three non-mutually exclusive mechanisms, namely overdominance, pleiotropy, and epistasis (Scheiner 1993). Of the three models, only overdominance made an explicit prediction about the relationship between plasticity and heterozygosity. The lack of strong evidence that heterozygous genotypes were more robust has led to the dismissal of the overdominance model, and current studies on the molecular basis of plasticity either consider the epigenetic and transcriptomic landscapes (*i.e.* the pleiotropy model) or the shape of regulatory networks (*i.e.* the epistasis model) (Goldstein and Ehrenreich 2021; Chevin et al. 2022). Our analysis of abdominal pigmentation in multiple Drosophila species supports an association between plasticity and heterozygosity but in a positive rather than negative way, hence revisiting the classical overdominance model. However, as indicated by Scheiner (1993), the three models are not mutually-exclusive, and we will illustrate how our results can agree with the different models depending on the species and the sex considered.

In its simplest form, a gene regulatory network consists of a transcription factor activating or suppressing an effector gene. Of the five monogenic cases studied here, the underlying locus involves an effector gene, *t*, in one species (*D. erecta*) (Yassin et al. 2016a), and a transcription factor, *pdm3*, in four species (*D. kikkawai*, *D. jambulina*, *D. burlai* and *D.* cf. *bocqueti*) (Yassin et al. 2016b; Fukutomi et al. 2025) (Figure 6). Loss-of-function regulatory mutations of *t* reduce plasticity in *D. erecta* recessive light homozygotes (Figure 2B), and only slightly affect the degree of dominance, *i.e.* plasticity of the functional allele is nearly codominant near the upper thermal limit when both dark heterozygotes and homozygotes become lighter (Figure 2B). Indeed, the *t* regulatory element which is involved in female color variation in both *D. erecta* and *D. melanogaster* (Bastide et al. 2013; Endler et al. 2016; Gibert et al. 2017a) partly affects pigmentation plasticity in the later species (Gibert et al. 2016). On the other hand, mutations in the transcription factor *pdm3* show more complex relationships (Figures 2B). First, the plasticity of both homozygotes shows almost similar slopes (but different means). Second, dominance relationship differed between species, even among sister species such as *D. burlai* and *D.* cf. *bocqueti*. In two species (*D. kikkawai* and *D. burlai*), the dark allele is dominant throughout the temperature range, whereas in the two other species (*D. jambulina* and *D.* cf. *bocqueti*), a strong dominance reversal was observed in females. Polymorphism in *pdm3* associates with female abdominal pigmentation variation in European and African populations of *D. melanogaster* (Endler et al. 2016). Mutations in this gene also underlie pigmentation evolution between the sister species *D. yakuba* and *D. santomea* with contrasting melanism (Liu et al. 2019; David et al. 2022). Whereas pigmentation plasticity in these two species is low, their reciprocal hybrids, which are heterozygous for *pdm3* interspecific differences along with other genes, are highly plastic (Figure 2A; Figure 5B). This may indicate that the relation between heterozygosity and plasticity may differ according to the type of the underlying gene.

**Figure 6.**
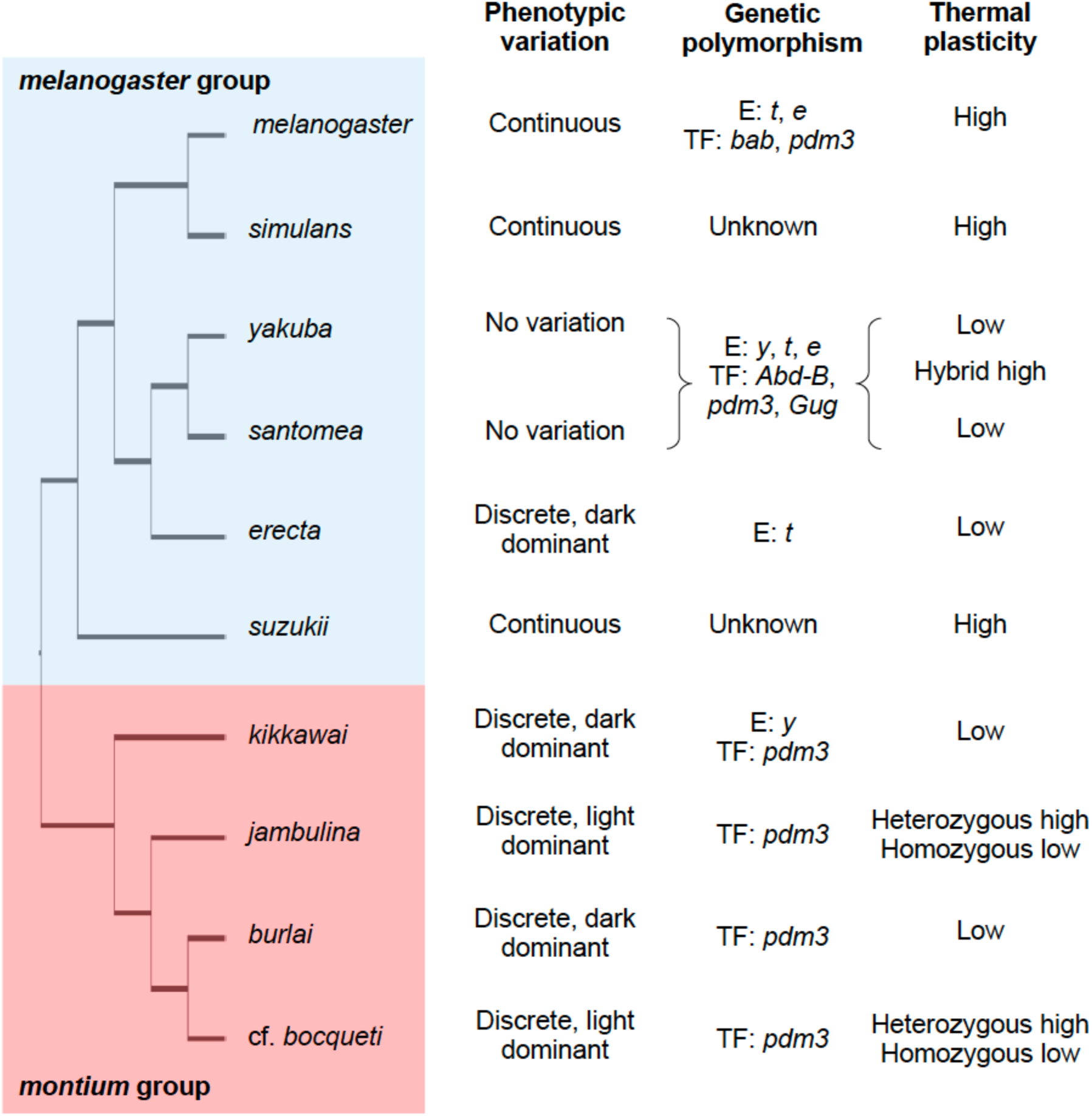
Evolution of the genetic architecture of pigmentation variation and its thermal plasticity in the *melanogaster* and *montium* species groups. E = effector melanin-synthesis gene; TF = transcription factor. Phylogenetic relationships are after Suvorov et al. (2022).

Besides, the types of the mutations in the alleles of the polymorphic gene may also affect the degree of the plastic response. Indeed, plasticity was highest in the heterozygotes of the two *montium* group species, *D. jambulina* and *D.* cf. *bocqueti*, where the light allele was dominant under optimal developmental temperature regimes, *i.e.* ≥21°C. The reversal of dominance according to the temperature, the main reason of the steepness of the reaction norm, is associated with the evolution of dominance between the *pdm3* alleles in these species. In *D. melanogaster*, the wildtype light allele is dominant (Tichy et al. 2008), but RNAi lines targeting different parts of this gene yields contrasting results, either increasing pigmentation (mostly in females) or decreasing it (mostly in males) (Rogers et al. 2014). Consequently, it is possible that the *pdm3* mutations underlying the dark allele in *montium* species where it is dominant, such as in *D. kikkawai* and *D. burlai*, are of the dominant negative type, which are common for transcription factors (Gerasimavicius et al. 2022). Such transcription factors usually have multiple isoforms, one of them has the DNA binding domain but lacks polymerase or other protein binding domains that are needed to express a gene, whereas another isoform has both DNA and protein binding domains. Consequently, the isoforms have opposite effects on the trait. In the heterozygote, the isoform lacking the protein binding domain competes with the isoform bearing this domain, hence altering its function. On the other hand, mutations in species with the light allele being dominant, like *D. jambulina* and *D. cf. bocqueti*, may be of the classical loss-of-function type that may reduce but do not completely eliminate the effect of the wild-type light allele. Further molecular dissection of the underlying alleles is required to understand the evolution of dominance reversal at *pdm3*.

The effect of mutations in both modifier and effector genes, *i.e.* the epistatic model of plasticity, is best illustrated in *D. kikkawai* and the hybrids between *D. santomea* and *D. yakuba*. In *D. kikkawai*, the effector *yellow (y)* gene lacks a conserved *cis*-regulatory element leading to the loss of body pigmentation (Jeong et al. 2008). Because this gene is X-linked, a dosage effect may occur between the sexes. Plasticity in males additively increased with the number of copies of the dominant dark *pdm3* allele, which is an indirect regulator of *y* (Liu et al. 2019). Putatively negative dominant mutations in this gene are expected to activate *y*, but the effect may depend on the number of copies of *y*, being higher in females. Similarly, whereas hybrid males mothered by *D. santomea*, *i.e.* those carrying defective X-linked melanin synthesis genes *t* and *y*, had similar plasticity slopes to hybrid females. But this slope was halved in hybrid males mothered by *D. yakuba*, with functional *y* and *t*. Therefore, plasticity is affected by the functionality of the polymorphism at multiple genes, as predicted by the epistasis model, although the degree of plasticity was higher in fully heterozygous background.

### The association between plasticity and polymorphism may be more general than currently appreciated

We have restricted our analyses to eight species belonging to two species groups of the subgenus *Sophophora*. Despite the differences in their reaction norms, both groups contain species with sexually dimorphic melanism, and a single gene with antagonistic sexual effect may underlie the association between plasticity and heterozygosity, hence reducing the independence of our observations. Future analyses should investigate pigmentation thermal plasticity in non-*Sophophora* fly species lacking sexual dimorphism, with or without Mendelian polymorphism (e.g., Brisson et al. 2005; Rocha et al. 2009; Wittkopp et al. 2009). Beyond *Drosophila*, color polymorphism abounds in nature (Orteu and Jiggins 2020; Sapir et al. 2021). Investigation of the diversity in reaction norms in species with heterozygous and homozygous color genotypes, especially in these with a hierarchy of dominance (e.g., Durand et al. 2014; Le Poul et al. 2014), should shed light on the generality of the relationship between heterozygosity and the degree of plasticity.

Most quantitative traits for which phenotypic plasticity was studied in *Drosophila*, such as body size, wing shape, bristle number, ovariole number or life-history traits, are highly polygenic with often a redundant genetic basis (David et al. 2004; Gibert et al. 2004a; Mackay and Lyman 2005; Lobell et al. 2017; Lafuente et al. 2018; Pitchers et al. 2019). Identifying the underlying genotypes of these traits to test for the effect of heterozygosity may be difficult. An alternative to the ‘forward genetics’ approach that we leveraged here, is to use a ‘reverse genetics’ analysis, wherein the transcription level of polymorphic genes can be analyzed in homozygous vs. heterozygous states in different environments, even in the lack of knowledge about the phenotypic consequences of the genotypes. One such significant example comes from a study by Chen et al. (2015) who showed that 62% of genes with thermal plastic response in *D. melanogaster* (*i.e.* ∼51% of total genes) swapped allelic dominance in heterozygotes between cold and hot environments. Future similar transcriptomic analyses are therefore highly needed to test the generality of these patterns, especially these using hybrids between species.

### The evolutionary consequences of links between phenotypic plasticity and polymorphism

A link between phenotypic plasticity and polymorphism can provide simple explanations on how plasticity evolves. In fact, plasticity can be lost through any mechanism reducing genetic heterozygosity such as inbreeding or directional selection. Indeed, the ‘flexible stem hypothesis’ postulates that from a plastic ancestor, descendants with fixed phenotypes from the ancestral phenotypic spectrum can evolve (West-Eberhard 2003; Gibert 2017). A likely example from our data is the case of *D. santomea* and *D. yakuba*. According to the phylogeny given in Figure 6, it is likely that the ancestor of these species was highly plastic given the high plasticity of *D. melanogaster* (Gibert et al. 2000) and *D. suzukii* (Colinet and Kustre 2025). Pigmentation difference between *D. yakuba* and *D. santomea* is due to mutations in at least six loci (Liu et al. 2019; David et al. 2022), but it is unlikely that all these mutations have arisen simultaneously. This means that a substantial degree of heterozygosity segregated in the ancestor of these species, and consequently this ancestor has gradually lost its plasticity as more and more alleles became fixed in each lineage. Indeed, pigmentation plays a role in the reproductive isolation between these two species (David et al. 2022), and a hybrid zone on the African island of Sao Tomé between these two species exists (Lachaise et al. 2000; Llopart et al. 2005; Turissini and Matute 2017). A plastic hybrid is disadvantaged and the ‘flexible stem hypothesis’ may in essence reflects a simple model of ‘underdominance’ of the heterozygote.

Conversely, phenotypic plasticity can be gained by any mechanism that increases genetic variability and heterozygosity in a population, such as mutations, outbreeding (e.g., gene flow and introgression), and balancing selection. Indeed, any new mutation in a di- or polyploid organism first arises in the heterozygous state, and it has been argued that a heterozygous advantage is essential for the spread of new mutations and counteracting the loss by random genetic drift (Sellis et al. 2011). In cases where plasticity is adaptive, the higher plasticity of the heterozygote can facilitate the spread of this mutation. Such adaptive plasticity gain model is in essence an ‘overdominance’ model. An important consequence of the overdominance model is that plasticity could be a mechanism to maintain genetic diversity, *i.e.* rather than ‘buying time’ for genetic adaptation to occur, plasticity may be a major source for ‘saving time’ by maintaining standing genetic variation, such as in the case of introduced species or multivoltine species inhabiting a heterogeneous spatial or seasonal environment (Gulisija et al. 2016; Wittmann et al. 2017; Promy et al. 2023). Further theoretical and empirical investigations of the link between heterozygosity and plasticity will definitively improve our understanding of the evolution of both.

## Authors contribution

J.R.D. – conceptualization, formal analysis, investigation, resources; B.D. – resources, writing review and editing; P.F. – data curation, formal analysis, visualization; A.L. – data curation, formal analysis, visualization; A.D. – investigation, resources; S.M. – investigation, resources; P.G. – writing review and editing; A.Y. – conceptualization, data curation, methodology, supervision, visualization, writing original draft.

## Conflicts of interest

The authors declare no conflict of interest.

## Data availability statement

All data associated with this study are provided in a Figshare Folder and will be released upon publication.

## Acknowledgments

The authors thank Artyom Kopp for comments on the manuscript and for sharing with Yuichi Fukutomi the association between female-limited color dimorphism and polymorphism at the *pdm3* locus in *D. jambulina* and *D.* cf. *bocqueti*.

## Notes

### Competing Interest Statement

The authors have declared no competing interest.

### Summary of Updates

Materials and Methods extended; additional statistical analyses conducted; Figures 1-5 revised; author affiliations updated; two new Tables 2 and 3.

